# CD56 expression of intravascular trophoblasts defines a class of vasculopathy in preeclampsia and other pregnancy complications

**DOI:** 10.1101/293027

**Authors:** Peilin Zhang

**Author notes:** Correspondence: Peilin Zhang, MD., Ph.D Department of Pathology NYP-BMH 506 6^th^ ST. Brooklyn, NY 11215.

## Abstract

We have discovered the expression of CD56 on the intravascular trophoblasts in the maternal spiral artery at implantation sites, and we have also found the similar phenotypic switch of intravascular trophoblasts in the decidual vasculopathy in preeclampsia. Currently we have examined 124 placentas from the patients with preeclampsia and 84 placentas from patients with other pregnancy associated complications without preeclampsia. CD56 expression on the intravascular trophoblasts can be seen in classic decidual vasculopathy such as fibrinoid medial necrosis and acute atherosis. In addition, partial involvement of the decidual vessels with classic vasculopathy can also be identified by CD56 expression. The cellular components of the classic vasculopathy in preeclampsia showed immunoreactivity to cytokeratin and CD68, in addition to CD56 expression, indicating the fetal trophoblastic cell origin. The classic decidual vasculopathy including acute atherosis and fibrinoid medial necrosis can be unified and designated as CD56-related vasculopathy. The CD56 related vasculopathy is associated not only to preeclampsia but also to other pregnancy related complications, such as gestational diabetes, placental infarcts, intervillous thrombosis and other fetal distress syndromes. Our study defined a spectrum of decidual vasculopathy spanning from the classic preeclampsia to other complications important to pregnancy that can be highlighted by CD56 expression, pointing to different direction of pathogenesis of preeclampsia and other pregnancy associated complications.

## Introduction

Decidual (Maternal) vasculopathy is commonly associated with pregnancy induced hypertension or preeclampsia. There are three types of decidual vasculopathy that are clinically recognized, namely acute atherosis, fibrinoid medial necrosis and mural arterial hypertrophy (1). These types of vasculopathy are present in approximately 20-50% placentas from preeclampsia (1, 2). Two separate issues are commonly seen. On the one hand, no specific morphological features of decidual vessels are identified in the remaining 50-80% preeclamptic placentas with clinical manifestations (3, 4); on the other, there are morphological features of vasculopathy in normotensive patients with other pregnancy associated complications such as gestational diabetes, intervillous thrombosis, placenta infarcts, fetal meconium passage and aspiration, and occasionally normal term pregnancy without identifiable complications. How the vasculopathy occurs in preeclampsia and how to bridge these gaps in understanding the disease process remain poorly understood. Traditionally acute atherosis is characterized by the presence of foamy macrophages (cholesterol/lipid laden macrophages) within the intima of the maternal spiral artery in the similar fashion to aortic atherosclerosis, and these foamy macrophages are also phenotypically expressing CD68 (5–10). Fibrinoid medial necrosis is characterized by the fibrin-like eosinophilic hyalinized material within the maternal vascular walls immunohistochemically reactive to immunoglobulins and complements (5–10). Mural arterial hypertrophy is defined as the thickened decidual arteries similar to those in other tissue without trophoblastic remodeling and this type of vasculopathy is known to be more commonly associated with chronic hypertension (11, 12). Through extensive laboratory and animal studies, preeclampsia is thought to be caused by poor cytotrophoblastic remodeling of spiral artery in early implantation and subsequently poor angiogenesis of placental vascular development (13–16). We have discovered that the trophoblastic remodeling of decidual spiral artery in early implantation is associated with phenotypic switch of trophoblasts to be immunohistochemically reactive to CD56, a maternal antigen likely from the natural killer cells within the decidua, and these trophoblasts with CD56 expression persist in vasculopathy in preeclamptic placenta but not in normal term pregnancy. We expanded our study to include more patients and other pregnancy complications associated with these types of vasculopathy. Our data showed acute atherosis and fibrinoid medial necrosis represent the spectrum of morphologic changes in the same category of vasculopathy. Both of which are CD56-related vasculopathy not only seen in preeclampsia but other complications in late pregnancy.

## Material and methods

Prospectively 81 implantation sites (specimens including missed abortion, incomplete abortion, spontaneous abortion and abortions of medical reasons), 124 placentas with clinical indication of preeclampsia, and 84 placentas submitted for pathology examination for other pregnancy complications but no indication of preeclampsia for eight months prior to the study were collected and analyzed for decidual vascular changes with immunohistochemical staining for CD56, CD68, and cytokeratin (AE1/AE3). Pathologic changes, placental measurements and the associated clinical information and clinical diagnosis were collected at the time of clinical examination. Paraffin embedded tissues from the routine surgical pathology specimens and the routine hematoxylin & eosin stained pathology slides were examined using light microscopy under the circumstance of normal routine pathology service. No special procedures are employed. Implantation sites, maternal vasculopathy such acute atherosis and fibrinoid medial necrosis in preeclampsia or unknown clinical conditions were identified in routine clinical service, and subsequently examined by immunostaining for CD56 expression.

Immunohistochemical staining procedures are performed on paraffin embedded tissues using Leica Biosystems Bond III automated immunostaining system following the manufacturing instruction. CD56 monoclonal antibody was purchased for clinical in vitro diagnostics from Agilent DAKO (mouse monoclonal antibody against human, Cat. # M730401-2) with appropriate dilutions and controls. AE1/AE3 and CD68 monoclonal antibodies were from Agilent DAKO under catalogue numbers M351501-2 and M081401-2 (Agilent DAKO, CA).

## Results

### 1. Trophoblastic expression of CD56 in maternal spiral artery at implantation site

We have reviewed 81 implantation sites from the routine pathology practice, and the decidual vessels were examined by light microscopy. Trophoblastic remodeling of the maternal spiral arteries are easily identified in the background of decidua with extravillous trophoblasts. The vascular walls within the implantation sites show various degrees of fibrinoid medial necrosis, a typical morphological feature of decidual vasculopathy in preeclampsia (Figure 1). The morphologic features of acute atherosis are also seen occasionally, but much less common than those of fibrinoid medial necrosis. Mural arterial hypertrophy of decidual vessels are commonly present in the implantation sites but its clinical significance is uncertain given the early stages of morphologic changes in the decidua and implantation. It remains possible that the spiral artery remodeling is yet to happen to these decidual vessels with mural hypertrophy in early implantation. There are scattered lymphocytes within the decidua tissue. Immunohistochemical staining for CD56 expression showed strong membrane/cytoplasmic staining signals only on the intravascular trophoblasts, but not on extravillous trophoblasts in the decidua (Figure 1). These intravascular trophoblasts are also reactive to AE1/AE3, indicating the fetal trophoblastic origin. These intravascular trophoblasts are also weakly reactive to CD68, a macrophage/monocyte/histiocyte marker. The lymphocytes within the decidua are positive for CD56 expression, consistent with the uterine natural killer (NK) cells as expected since CD56 is a defining marker for uterine NK cells. The extravillous trophoblasts within the decidua outside vessels are reactive to both AE1/AE3 and CD68, but not to CD56. Prospectively, immunostaining for CD56 expression was performed for 30 consecutive cases (implantation sites), and 100% of the intravascular (intraluminal) trophoblasts were stained positive. It is apparent that the intravascular (intraluminal) trophoblasts have undergone the phenotypic switch by acquisition of CD56 expression in implantation sites, and there is no need for immunostaining for additional cases for confirmation.

**Figure 1:**
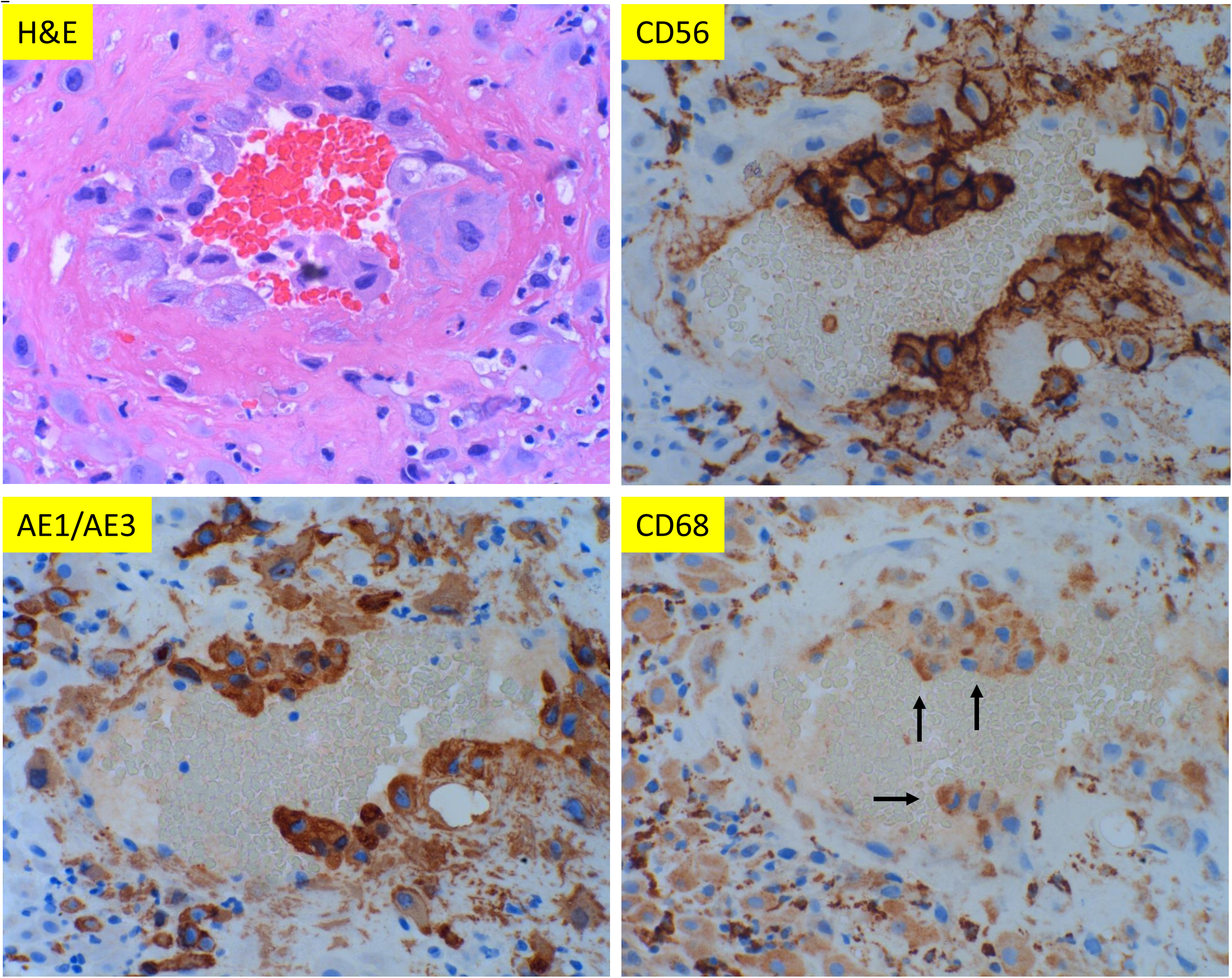
Morphologic features of maternal spiral artery remodeling in early implantation site with immunohistochemical staining for CD56, pancytokeratin (AE1/AE3) and CD68 expression. A: Implantation site of early missed abortion with hematoxylin & eosin (H & E) stain. B, C and D: The same section of implantation site as panel A with CD56, AE1/AE3 and CD68 immunostaining. (A-D at 400 X magnification). Arrow indicates weak CD68 reactivity.

### 2. Trophoblastic expression of CD56 in decidual vasculopathy in preeclampsia

Decidual vasculopathy including acute atherosis and fibrinoid medial necrosis is commonly present in the placentas from the patients with preeclampsia. Our previous survey indicated a relatively low percentage of placentas from the preeclamptic patients showed the presence of decidual vasculopathy (4, 17). Others also showed similar results (1, 18). Currently, we have examined 124 placentas from patients with clinical diagnosis of preeclampsia with additional knowledge of phenotypic switch of intravascular trophoblasts with CD56 expression above. The morphological features of decidual vasculopathy and the corresponding immunostaining for CD56, AE1/AE3 and CD68 expression are shown in Figure 2. The intravascular trophoblasts are again immunohistochemically reactive strongly to CD56, and these intravascular trophoblasts are also positive for AE1/AE3 and weakly positive for CD68. The presence of AE1/AE3 expression on these intravascular cells indicates the fetal trophoblastic cell origin, rather than maternal macrophages. This is in contrast to the traditional view of acute atherosis being similar to atherosclerosis with foamy macrophages (6, 8). The presence of CD68 expression on these intravascular trophoblasts, in addition to CD56 expression, may have other clinical implications, as CD68 expression is found in all syncytiotrophoblasts of the entire term placenta (data not shown). It is important to note that the decidual vessels with intravascular trophoblasts may or may not show complete fibrinoid medial necrosis or atherosis of the entire vessel wall. When fibrinoid medial necrosis replaces the entire walls of the vessels, there will be no cellular components, and no immunoreactive cells will be found by immunostaining. It is this partial replacement of the vascular wall with the residual cellular components that can be identified by the morphologic assessment and immunostaining. Four additional cases of placentas from the preeclamptic patients with corresponding CD56 immunostains are shown in Figure 3. This type of partial fibrinoid medial necrosis of vascular wall with intravascular cell components are not previously appreciated and recognized as a type of decidual vasculopathy (17). It is also important to note that the intravascular trophoblasts in preeclampsia were greatly reduced in number in comparison to those in the implantation sites (Figure 1), and the placentas from the normal term pregnancy show no evidence of intravascular trophoblasts nor typical vasculopathy (atherosis and fibrinoid medial necrosis). Acute atherosis and fibrinoid medial necrosis are frequently co-exist in the same vessels, and for practical purpose, these are designated as “classic” decidual vasculopathy. Mural arterial hypertrophy commonly associated with chronic hypertension and/or preeclampsia was not examined by the immunohistochemical staining for CD56 expression, since there is no intravascular trophoblast or CD56-positive cells (data not shown).

**Figure 2:**
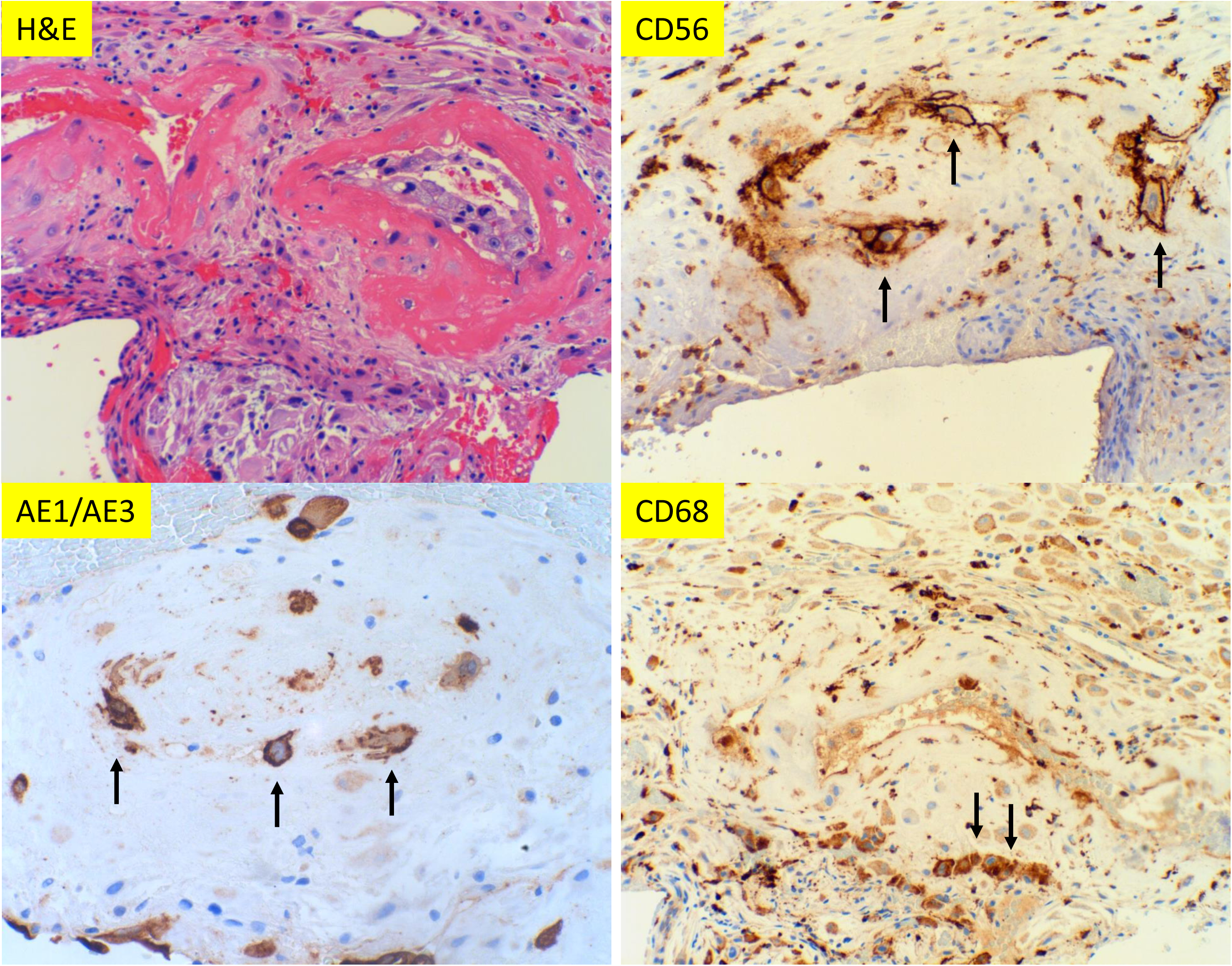
Morphologic features of classic decidual vasculopathy of preeclamptic placentas with immunohistochemical staining for CD56, AE1/AE3 and CD68 expressions. A: Classic decidual vasculopathy with acute atherosis and fibrinoid medial necrosis by hematoxylin & eosin (H & E) stain. B, C and D: The same section of placenta as panel A with CD56, AE1/AE3 and CD68 immunostaining. (A-D at 400 X magnification). Arrows indicate positive reactivity to individual markers.

**Figure 3:**
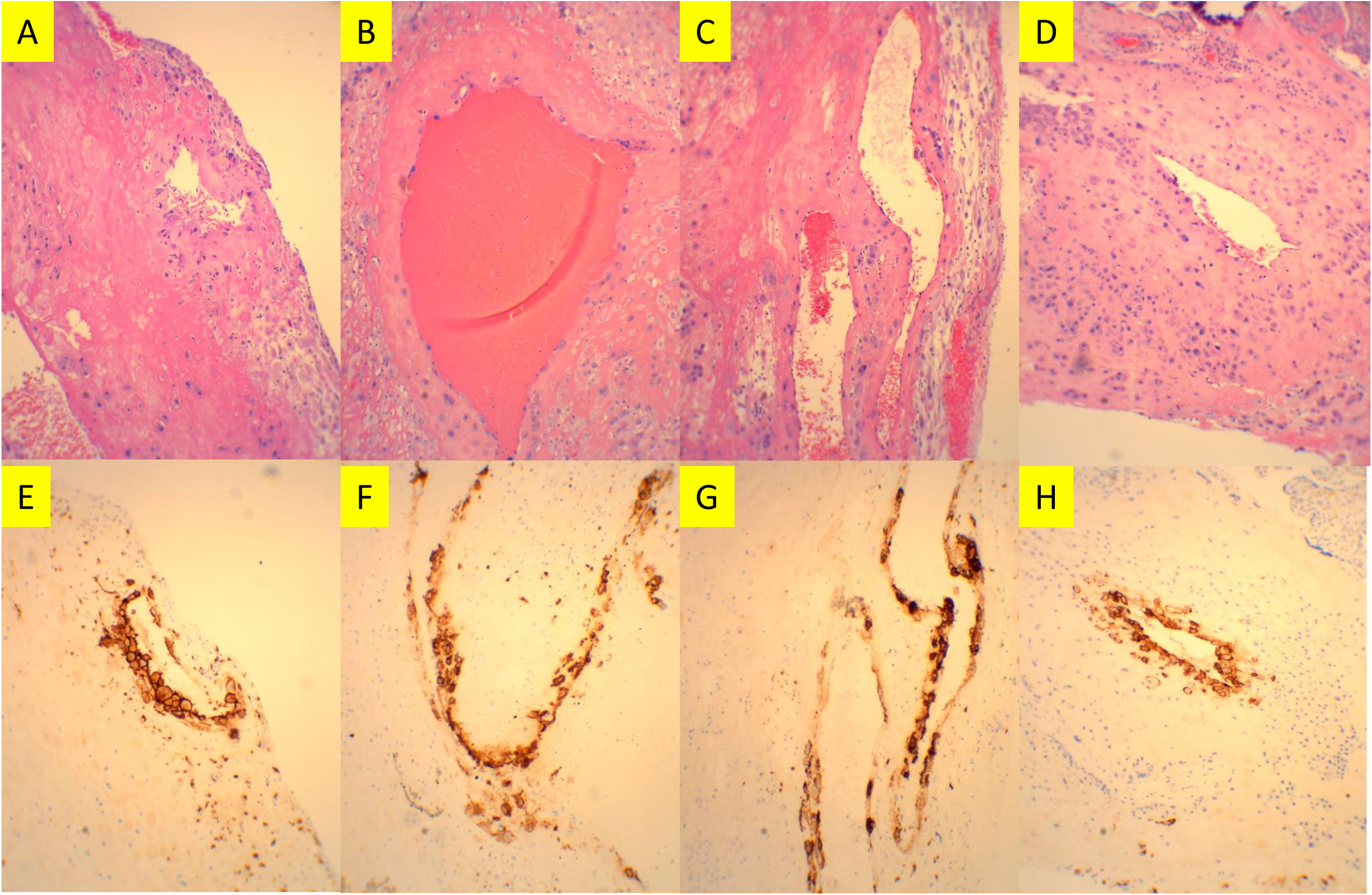
Four separate preeclamptic placentas with partial involvement of classic decidual vasculopathy and corresponding CD56 immunostaining patterns (all sections are at 100X magnification).

The clinical features of the 124 preeclamptic placentas are summarized in Table 1 based on the types of decidual vasculopathy and clinical/pathological complications. There were totally 78 (63%) preeclamptic placentas with classic decidual vasculopathy and 36 (29%) preeclamptic placentas with no evidence of decidual vasculopathy. The placental weight and the gestational age distributions of these preeclamptic placentas with and without vasculopathy are shown in Figure 4. The average (mean) weight for the preeclamptic placentas with classic vasculopathy is 438 grams, and for the preeclamptic placentas without vasculopathy 495 grams (p=0.02, unpaired student t-test) (Table 1). The average preeclamptic placental weight with decidual vasculopathy is statistically lower than that without vasculopathy. The average gestational ages of the placentas with or without vasculopathy showed no significant difference. Immunostains for CD56 was performed for 39 cases, and 37 cases showed demonstrable immunoreactivity to CD56 (95%). Two negative cases for CD56 immunostaining were due to the disappearance of the decidual vessels on the immunostaining slides and the deeper sections. The most common placental abnormality associated with classic decidual vasculopathy is infarcts (22%) followed by intervillous thrombosis (17%). Meconium stain of fetal membrane, fetal surface or amniotic fluid (15%), gestational diabetes (10%), and inflammatory conditions (9%) are also commonly associated with the presence of classic decidual vasculopathy. Intrauterine growth restriction (IUGR), category 2 fetal heart tracing during labor, placental abruption (microscopic) and intrauterine fetal demise (IUFD) are present less frequently. There are 22% preeclamptic placentas with decidual vasculopathy showing no additional identifiable placental abnormalities.

**Table 1:**
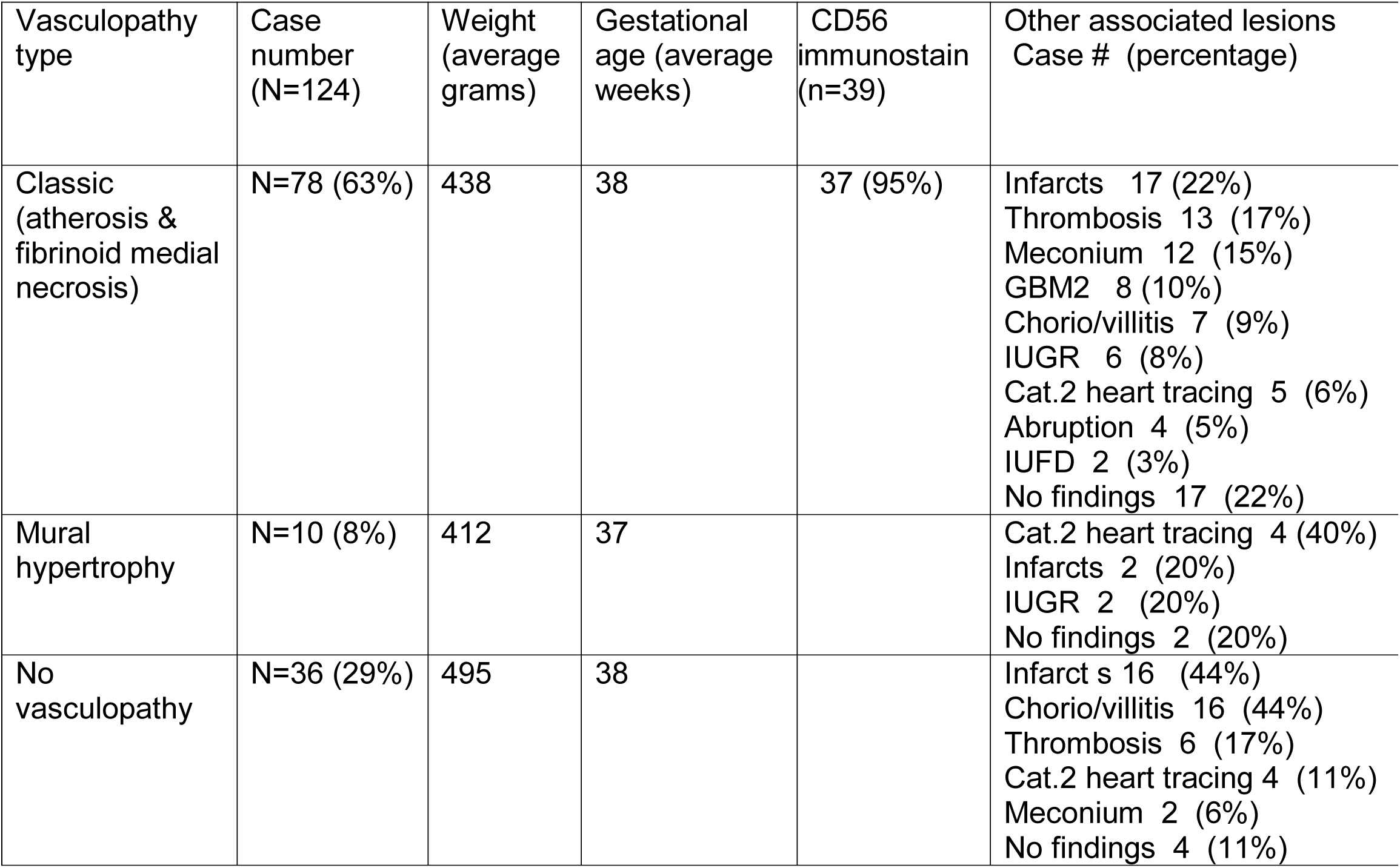
Preeclamptic placentas with decidual vasculopathy and complications

**Figure 4:**
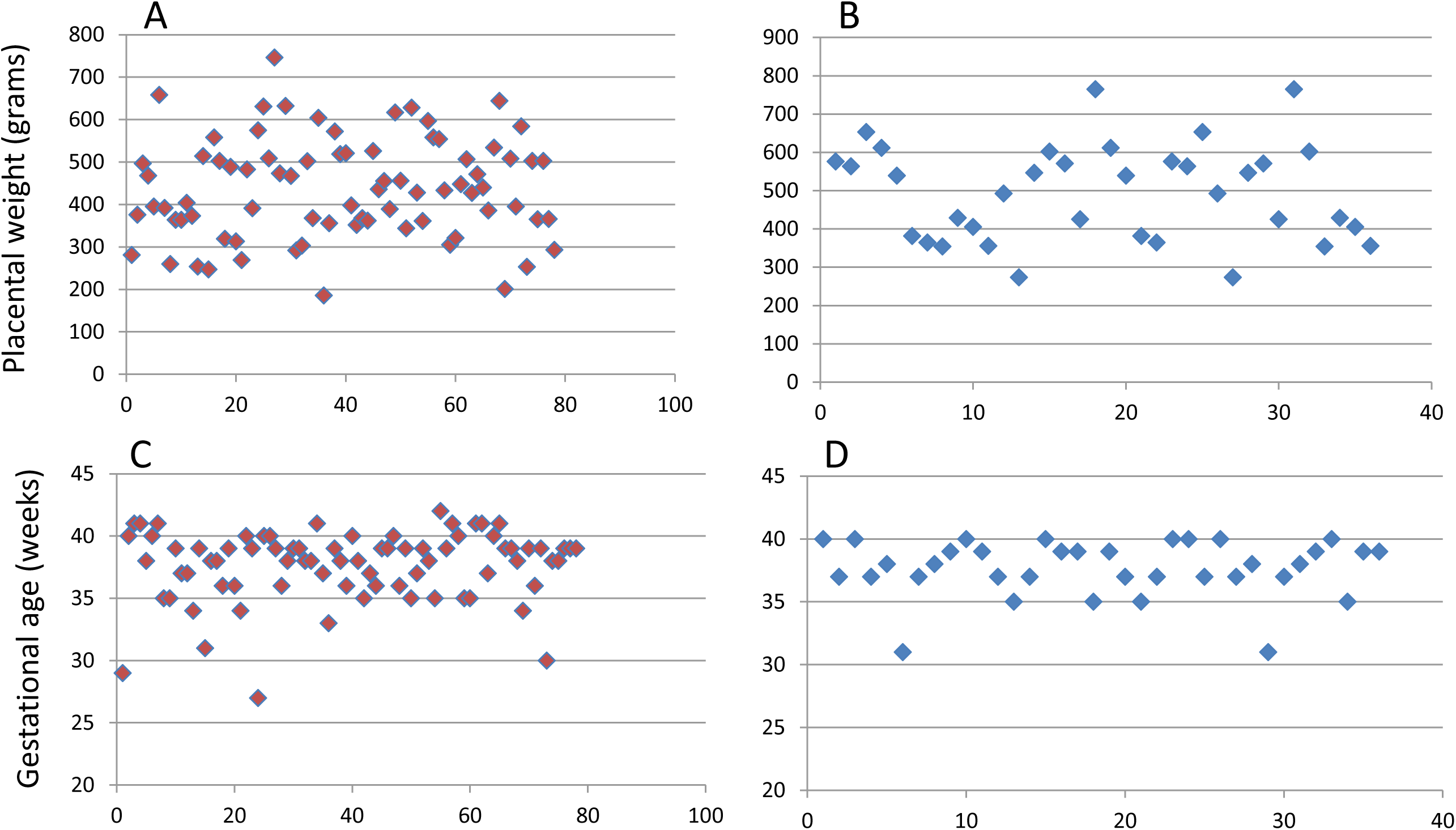
Placental weight and gestational age distributions between the 78 preeclamptic placentas with decidual vasculopathy and 36 preeclamptic placentas without decidual vasculopathy. The placental weight is statistically different (p=0.02, unpaired student t-test), and the gestational age is not. A and B represent the preeclamptic placental weights with (n=78) or without (n=36) decidual vasculopathy. C and D represents the gestational ages of the preeclamptic placentas with or without decidual vasculopathy. X-axis represents individual case number.

The preeclamptic placentas without classic decidual vasculopathy (totally n=36) are also associated with other significant pathologic abnormalities such as infarcts (44%), inflammatory conditions (44%), intervillous thrombosis (17%), category 2 heart tracing during labor (11%), and meconium passage (6%). There were 11% preeclamptic placentas with no decidual vasculopathy showing no additional pathologic abnormalities.

### 3. Decidual vasculopathy without preeclampsia

There are clinical scenarios with complications in pregnancy such gestational diabetes, maternal thrombotic conditions, maternal fever, or abnormal fetal heart rate tracing during labor without co-existing diagnosis of preeclampsia. We have examined 84 consecutive placental cases with decidual vasculopathy and other pregnancy related complications, and these patients did not have diagnosis of preeclampsia or hypertension. The pathologic findings of these placentas are shown in Table 2. The morphological features of decidual vasculopathy were identical to those seen in Figure 2. The average (mean) placental weight from these non-preeclamptic patients (464 grams) shows no statistical difference from that of the preeclamptic patients (448 grams) (p=0.9, unpaired student t-test). The placental weight and the gestational age distributions of the 124 preeclamptic placentas and the 84 non-preeclamptic placentas are shown in Figure 5. Immunostaining for CD56 expression were performed on 28 cases with identifiable decidual vasculopathy and all 28 cases were positive for CD56 reactivity (100%). The most common associated placental abnormalities from these non-preeclamptic placentas are similar to those seen in preeclamptic patients including infarcts (11%), thrombosis (17%), category 2 fetal heart tracing during labor (20%), meconium passages (23%), and inflammatory conditions such as chorioamnionitis or villitis (15%) (Table 2) with less frequent complications including IUGR (5%), abruption (4%) and IUFD (1%). There were 14% non-preeclamptic placentas with decidual vasculopathy showing no additional pathologic findings.

**Table 2:** Placentas with decidual vasculopathy but not preeclampsia

| Case # | Average weight (grams) | Gestational age (weeks) | CD56 immunostain (n=28) | Associated lesions N=84 |
| --- | --- | --- | --- | --- |
| N=84 | 464 | 39 | 28 (100%) | Meconium 19 (23%)<br>Cat. 2 heart tracing 17 (20%)<br>Thrombosis 14 (17%)<br>Chorio/villitis 13 (15%)<br>GBM1/2 10 (12%)<br>Infarcts 9 (11%)<br>IUGR 4 (5%)<br>Abruptio 3 (4%)<br>IUFD 1 (1%)<br>Cholestasis 1 (1%)<br>No other findings 12 (14%) |

**Figure 5:**
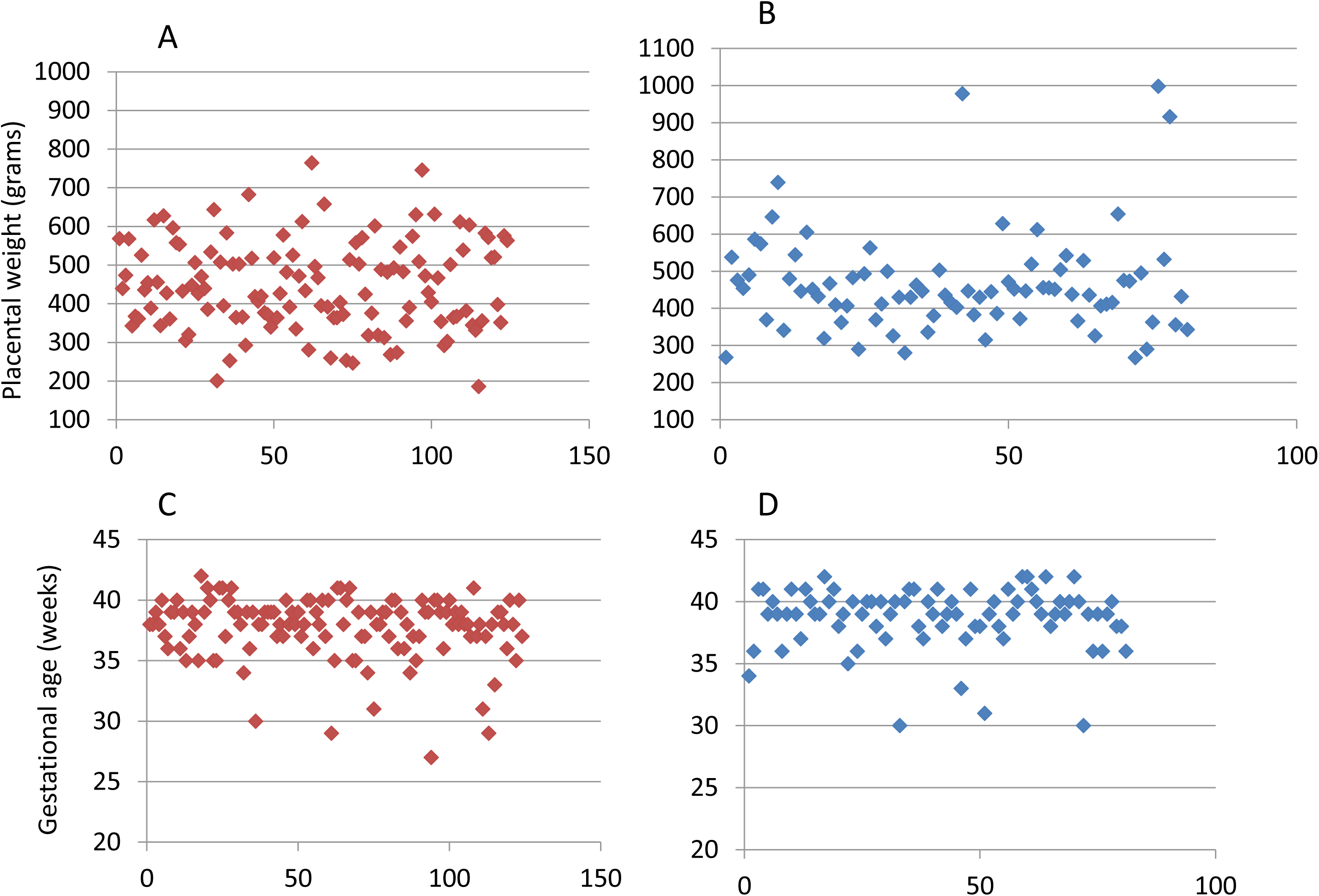
The placental weight and gestational age distributions of the 124 preeclamptic placentas and 84 non-preeclamptic placentas with decidual vasculopathy. The placental weight and gestational age show no statistically significant difference. A and B represent the weight distributions of the preeclamptic (n=124) and non-preeclamptic (n=84) placentas. C and D represent the gestational age distribution of preeclamptic (n=124) and non-preeclamptic (n=84) placentas.

## Discussion

Preeclampsia is a spectrum of clinical manifestations with various degrees of severities and the placental manifestations of preeclampsia are also diverse. Decidual vasculopathy is one of the defining features of preeclamptic placentas. In severe preeclampsia which represents approximately 5-20% of all preeclamptic patients, the placental villous tissue is hypoplastic with commonly associated decidual vasculopathy, placental infarcts and thrombosis (19). In these cases, the pathological changes of the placentas are easily identified on routine placental examination. Much commonly seen are mild forms of preeclampsia with borderline normal placentas and normal villous tissue development. In this mild form of preeclamptic cases, the placental pathological changes are harder to define, and decidual vasculopathy difficult to identify (17, 19). CD56 immunostaining can be used as an adjunct marker for identifying the intravascular (intraluminal) trophoblasts, and decidual vasculopathy. The spectrum of decidual vasculopathy from typical acute atherosis, fibrinoid medial necrosis to partially involved decidual vessels by these changes can be defined by the immunostaining for CD56 expression. Practically these decidual vascular lesions in preeclampsia are probably better designated as “CD56 related vasculopathy”, in comparison to another vascular lesion defined as “mural arterial hypertrophy”. In addition, the term “CD56 related vasculopathy” appears to unify the spectrum of morphologic changes in decidual vasculopathy with implication of pathogenesis. Mural arterial hypertrophy is shown to be associated with significant pregnancy related complications (11). In light of CD56 expression of intravascular trophoblasts in classic decidual vasculopathy in this study, mural arterial hypertrophy seems to be a separate vascular entity with different pathogenic mechanism. Interestingly, CD56 expression is only found on intravascular trophoblasts and the scattered decidual NK cells. No other cell type in early implantation site or term placenta is found to be immunoreactive to CD56 expression. Traditionally acute atherosis and fibrinoid medial necrosis are related morphologic changes characterized by the endothelial cell damage, immunoglobulins/complements deposits, fat/cholesterol deposits and macrophage phagocytosis (5–7, 10, 20). Our current results demonstrated that these “macrophages” are phenotypically fetal trophoblasts in origin by immunostaining pattern for cytokeratin (AE1/AE3) expression. These “macrophages” in the vessels are also immunoreactive to CD56 expression and weakly reactive to CD68 expression. Identification of these “foamy macrophages” as fetal trophoblasts with expression of CD56, a defining NK cells marker, points to an entirely different direction for pathogenesis of preeclampsia. CD56 expression is present in all implantation sites, and not present in the normal term placentas, indicating the disappearance of CD56 expression and/or intravascular trophoblasts is essential for normal pregnancy. Consequently, persistence of intravascular trophoblasts with CD56 expression at the late gestations is associated with pregnancy related complications. This is in contrast to the traditional view of the lack of spiral artery remodeling by the trophoblasts as a basis of preeclampsia (13–16). We have seen CD56 expression on intravascular trophoblasts in placentas of up to 24 week gestation from early premature delivery due to incompetent cervix or chorioamnionitis without evidence of decidual vascular insufficiency (maternal vascular malperfusion). Exactly how CD56 expression disappears in temporal fashion from early to late gestations requires further laboratory studies.

In light of immunostaining and knowledge of CD56 expression in decidual vasculopathy, there are still 29% preeclamptic placentas without identifiable decidual vasculopathy in the current study (Table 1). These preeclamptic placentas contain significant pathologic lesions such as infarcts, intervillous thrombosis and other findings similar to those identified in placentas with decidual vasculopathy. One of the possible reasons may relate to the placental sampling, since the routine placental examination requires three sections of placental tissue including maternal and fetal surfaces as recommended in the international guideline (17). Decidual vascular changes and vasculopathy are best appreciated on the decidua basalis (basal plate), and additional sampling in the basal plate may increase the yield of decidual vasculopathy.

Maternal spiral artery remodeling by the extravillous trophoblasts is critical for establishment of embryonic implantation in the endometrium (21). The expression of CD56 only on the intravascular (intraluminal) trophoblasts, but not on the decidua extravillous trophoblasts suggests a dynamic interaction between the fetal trophoblasts and the maternal uterine NK cells upon invasion into the vascular wall. It has been known that the maternal uterine NK cells can regulate the trophoblastic remodeling of maternal spiral artery by an unclear mechanism (22–24), and direct acquisition of maternal NK cell marker CD56 by the fetal trophoblasts seems to help explain the direct interaction between the maternal and fetal components. CD56 is a glycoprotein first discovered in neuron, glial and neural tissues (neural cell adhesion molecule, NCAM). CD56 is also expressed in the inner cell mass of early embryonic development (25). In hematopoietic cells, CD56 marks the NK cell lineage and minor percentage of T cells, and aberrant expression of CD56 can be found in multiple myeloma cells and acute myeloid leukemia (26). The role of the uterine NK cells in implantation appears to be a direct interaction as a provider of the maternal CD56 to the intravascular (intraluminal) trophoblasts, in addition to autocrine and paracrine functions providing cytokines and secreting factors to the fetal development. Direct acquisition of maternal CD56 expression by the intravascular trophoblasts may have significance to suppress the maternal immune system or to induce maternal immune tolerance since invasion of trophoblasts into the maternal spiral artery is the initial exposure of the fetal cells to the maternal circulation (27). The mechanism of the interaction between the fetal trophoblasts and the maternal NK cells should be further explored experimentally using in vitro models.

There is a soluble form of CD56 in the circulation (26, 28), and it will be interesting and useful to see if the levels of the soluble CD56 is associated with development of preeclampsia.

**Table 1:** Summary of the124 preeclamptic placentas with different types of decidual vasculopathy and additional pathologic findings.

**Table 2:** Summary of the 84 non-preeclamptic placentas with decidual vasculopathy and additional pathologic findings.

## Reference

1. Benirschke K, Burton, Graham J., Baergen, Rebecca N. Pathology of the Human Placenta. 2012.

2. Ananth CV, Keyes KM, Wapner RJ. Pre-eclampsia rates in the United States, 1980-2010: age-period-cohort analysis. Bmj. 2013;347:f6564.

3. Kim YM, Chaemsaithong P, Romero R, et al. Placental lesions associated with acute atherosis. J Matern Fetal Neonatal Med. 2015;28:1554–1562.

4. Zhang P, Schmidt M, Cook L. Maternal vasculopathy and histologic diagnosis of preeclampsia: poor correlation of histologic changes and clinical manifestation. Am J Obstet Gynecol. 2006;194:1050–1056.

5. Robertson WB, Brosens I, Dixon HG. The pathological response of the vessels of the placental bed to hypertensive pregnancy. J Pathol Bacteriol. 1967;93:581–592.

6. Labarrere C, Alonso J, Manni J, et al. Immunohistochemical findings in acute atherosis associated with intrauterine growth retardation. Am J Reprod Immunol Microbiol. 1985;7:149–155.

7. Labarrere CA. Acute atherosis. A histopathological hallmark of immune aggression? Placenta. 1988;9:95–108.

8. Kitzmiller JL, Watt N, Driscoll SG. Decidual arteriopathy in hypertension and diabetes in pregnancy: immunofluorescent studies. Am J Obstet Gynecol. 1981;141:773–779.

9. Kitzmiller JL, Aiello LM, Kaldany A, et al. Diabetic vascular disease complicating pregnancy. Clin Obstet Gynecol. 1981;24:107–123.

10. Kitzmiller JL, Benirschke K. Immunofluorescent study of placental bed vessels in pre-eclampsia of pregnancy. Am J Obstet Gynecol. 1973;115:248–251.

11. Labarrere CA, DiCarlo HL, Bammerlin E, et al. Failure of physiologic transformation of spiral arteries, endothelial and trophoblast cell activation, and acute atherosis in the basal plate of the placenta. Am J Obstet Gynecol. 2017;216:287 e281–287 e216.

12. Hecht JL, Zsengeller ZK, Spiel M, et al. Revisiting decidual vasculopathy. Placenta. 2016;42:37–43.

13. Norwitz ER, Schust DJ, Fisher SJ. Implantation and the survival of early pregnancy. N Engl J Med. 2001;345:1400–1408.

14. Cross JC, Werb Z, Fisher SJ. Implantation and the placenta: key pieces of the development puzzle. Science. 1994;266:1508–1518.

15. Goldman-Wohl DS, Ariel I, Greenfield C, et al. Lack of human leukocyte antigen-G expression in extravillous trophoblasts is associated with pre-eclampsia. Mol Hum Reprod. 2000;6:88–95.

16. Roberts JM, Taylor RN, Musci TJ, et al. Preeclampsia: an endothelial cell disorder. Am J Obstet Gynecol. 1989;161:1200–1204.

17. Khong TY, Mooney EE, Ariel I, et al. Sampling and Definitions of Placental Lesions: Amsterdam Placental Workshop Group Consensus Statement. Arch Pathol Lab Med. 2016;140:698–713.

18. Kim YM, Chaemsaithong P, Romero R, et al. The frequency of acute atherosis in normal pregnancy and preterm labor, preeclampsia, small-for-gestational age, fetal death and midtrimester spontaneous abortion. J Matern Fetal Neonatal Med. 2015;28:2001–2009.

19. Redline RW, Boyd T, Campbell V, et al. Maternal vascular underperfusion: nosology and reproducibility of placental reaction patterns. Pediatr Dev Pathol. 2004;7:237–249.

20. Brosens I, Robertson WB, Dixon HG. The physiological response of the vessels of the placental bed to normal pregnancy. J Pathol Bacteriol. 1967;93:569–579.

21. Red-Horse K, Zhou Y, Genbacev O, et al. Trophoblast differentiation during embryo implantation and formation of the maternal-fetal interface. J Clin Invest. 2004;114:744–754.

22. Lash GE, Otun HA, Innes BA, et al. Regulation of extravillous trophoblast invasion by uterine natural killer cells is dependent on gestational age. Hum Reprod. 2010;25:1137–1145.

23. Lash GE, Robson SC, Bulmer JN. Review: Functional role of uterine natural killer (uNK) cells in human early pregnancy decidua. Placenta. 2010;31 Suppl:S87–92.

24. Hanna J, Goldman-Wohl D, Hamani Y, et al. Decidual NK cells regulate key developmental processes at the human fetal-maternal interface. Nat Med. 2006;12:1065–1074.

25. Sundberg M, Jansson L, Ketolainen J, et al. CD marker expression profiles of human embryonic stem cells and their neural derivatives, determined using flow-cytometric analysis, reveal a novel CD marker for exclusion of pluripotent stem cells. Stem Cell Res. 2009;2:113–124.

26. Kaiser U, Oldenburg M, Jaques G, et al. Soluble CD56 (NCAM): a new differential-diagnostic and prognostic marker in multiple myeloma. Ann Hematol. 1996;73:121–126.

27. Fu B, Li X, Sun R, et al. Natural killer cells promote immune tolerance by regulating inflammatory TH17 cells at the human maternal-fetal interface. Proc Natl Acad Sci U S A. 2013;110:E231–240.

28. Taylor RN, Heilbron DC, Roberts JM. Growth factor activity in the blood of women in whom preeclampsia develops is elevated from early pregnancy. Am J Obstet Gynecol. 1990;163:1839–1844.

